# Statistical modelling of seafood fraud in the Canadian supply chain

**DOI:** 10.1101/2024.02.05.578947

**Authors:** Jarrett D. Phillips, Fynn A. De Vuono-Fraser

**Author notes:** **Corresponding Author**: Jarrett D. Phillips^1^ **Email Address**.

## Abstract

Seafood misrepresentation, encompassing product adulteration, mislabelling, and substitution, among other fraudulent practices, has been rising globally over the past decade, greatly impacting both the loss of important fish species and the behaviour of human consumers alike. While much effort has been spent attempting to localise the extent of seafood mislabelling within the supply chain, strong associations likely existing among key players have prevented timely management and swift action within Canada and the USA in comparison to European nations. To better address these shortcomings, herein frequentist and Bayesian logistic Generalised Linear Models (GLMs) are developed in R and Stan for estimation, prediction and classification of product mislabelling in Metro Vancouver, British Columbia, Canada. Obtained results based on odds ratios and probabilities paint a grim picture and are consistent with general trends found in past studies. This work paves the way to rapidly assess the current state of knowledge surrounding seafood fraud nationally and on a global scale using established statistical methodology.

## 1 Introduction

Seafood fraud is an escalating problem impacting society today, where both economic loss and substantial harm to fish populations have been sorely felt, especially in Canada and the USA (Cawthorn et al., 2021; Hanner et al., 2011; Hu et al., 2018; Naaum and Hanner, 2015; Naaum et al., 2015; Shehata et al., 2019, 2018; Warner et al., 2019). In its simplest form, seafood fraud is the practice of misleading consumers about products to increase profits (Naaum et al., 2016). Canada is not exempt in this respect, and is well above the global average mislabelling rate of 30% found by Pardo et al. (2016) in an examination of 51 published studies targeting 4500 samples. One aspect of food fraud pertaining especially to the Canadian market in comparison to countries within Europe, where specific labelling laws and up-to-date and comparable regulatory seafood lists are more tightly regulated (Cawthorn et al., 2021), is seafood product mislabelling. Unfortunately, Canada and the USA have a unique problem where labelling policies, gleaned through the Canadian Food Inspection Agency (CFIA) Fish List and the United States Food and Drug Administration (USFDA) Seafood List respectively, are extremely vague. This has led to the compounding of seafood mislabelling rates at all levels of the supply chain both outside of and within Canada (Shehata et al., 2019). For example, Canadian import guidelines allow over 200 species to be labelled and sold as snapper, a group which includes vulnerable species such as the Northern red snapper (*Lutjanus campechanus*). Typically, determination of species-level mislabelling is achieved on the basis of DNA-based specimen identification techniques such as DNA barcoding (Hebert et al., 2003a,b). Estimates suggest mislabelling rates of 50% and higher throughout the supply chain, but reliability largely depends on the maturity of genomic reference sequence databases, such as the Barcode of Life Data Systems (BOLD) (Ratnasingham and Hebert, 2007), upon which confident species-level matches can be readily ascertained from high-quality taxon records (Phillips et al., 2023). This, in turn, is strongly driven by levels of standing intraspecific haplotype variation and the presence of sufficiently wide DNA barcode gaps separating within- *versus* among-species marker diversity (Phillips et al., 2019, 2022). To ensure DNA sequence databases are populated with statistically defensible data, algorithms like HACSim (Phillips et al., 2020) should be employed to estimate required specimen sample sizes needed to capture desired levels of existing species genetic variation. Wong and Hanner (2008) were among the first to assess the efficacy of barcode reference libraries in identifying mislabelled seafood collected from markets across Canada and the USA. Out of the 91 collected samples that successfully amplified, 25% were found to be mislabelled (Wong and Hanner, 2008). Despite high fraud levels within the seafood supply chain, the spatiotemporal parameters of consumer product labelling remain obscure, but are not isolated to a single source (Shehata et al., 2019). Species mislabelling manifests at all times of year and involves many key participants within the supply chain hierarchy, including fishers and retailers. Strong correlations also likely exist with species stock abundance, market price, and conservation status, hindering reliable fraud prediction.

The extent to which seafood products are mislabelled depends on a number of factors, such as species biomass availability, market price, and conservation status. Further, product adulteration can be accidental and/or deliberate and can occur across different stages of seafood production, processing, and sale. Non-trivial differences in prices between wholesale- and retail-purchased seafood has increased opportunities for market fraud. Improving access to provenance history in the form of chain of custody documents will ensure transparency of seafood is not compromised. A widespread pattern in the seafood supply chain is the substitution of species of higher market value with those of lower market value in order to increase profitability by deceiving buyers and bypassing tariffs on product import. High-cost species are often of conservation concern due to decreasing population numbers as a result of overharvesting fueled by rising consumer demand, and thus are often substituted with highly-abundant low cost species (Zander and Feucht, 2018). Seafood mislabelling is also exercised as a means to conceal at-risk species, such as sharks, sourced from Illegal Unreported and Unregulated (IUU) fisheries for the sale of shark fins within the Asian market (Hanner et al., 2016). Instances of mislabelling can also pose serious health risks (Shehata et al., 2020) or challenge established religious doctrines and lifestyle choices. For instance, over-consumption of escolar (*Lepidocybium flavobrunneum*), which is often misrepresented as butterfish or white (Albacore) tuna, can lead to serious gastrointestinal issues along with other symptoms (Pardo et al., 2016). As a result, the sale of escolar is banned in many countries such as Italy, Japan, and South Korea. All the abovementioned behaviours seem to suggest that seafood fraud is intentional, as seafood product mislabelling in the Canadian economy appears to have risen in the past decade, negatively impacting the environmental sustainability of fish stocks. Oceana Canada, a not-for-profit charitable ocean conservation organisation subsidiary of the larger Oceana, has conducted numerous national surveys highlighting the detrimental impacts of seafood mislabelling across several major Canadian metropolitan cities in recent years. Findings are unsurprising: their 2018 survey indicated an overall mislabelling rate of 44% across nearly 400 samples and five large cities (Halifax, Ottawa, Toronto, Vancouver, and Victoria), with Victoria having the highest mislabelling average at 67%, and snapper being among the most frauded species (Oceana, 2018). This trend appears to be on the rise as evidenced from a more recent survey conducted in 2021 which saw a 47% mislabelling rate across 94 samples collected from four metropolitan cities (Halifax, Montreal, Ottawa, and Toronto) (Oceana, 2021).

While there have been hundreds of studies published on seafood fraud over the past decade, using basic statistical approaches, such as chi-square tests, the majority of research has simply demonstrated that mislabelling is evident and still largely persists within the supply chain on a global scale. Despite this, a handful of publications have attempted, with some degree of success, to quantify the effects of economically-motivated fraud through the quantification of overall mislabelling rates. Unfortunately, many do not report measures of uncertainty around estimates such as standard errors (SEs) or confidence intervals (CIs). Overall mislabelling rates can be assumed to approximately follow a Binomial distribution as individual seafood product samples are either correctly labelled, or not. That is, each collected sample can be considered an independent Bernoulli trial. Thus, a näıve way forward in this regard is to calculate the estimated Wald SE of the sample proportion of mislabelled samples, which is given by

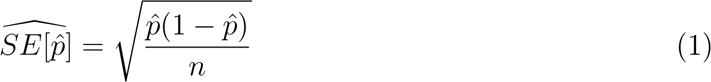

where 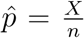 is the Maximum Likelihood Estimator (MLE) of the population mislabelling rate, *X* is the number of mislabelled observations, and *n* is the number of samples. However, the above formula is problematic for several reasons. First, Equation (1) is a Normal approximation based on large-sample theory through the Central Limit Theorem (CLT); thus, large sample sizes are required for reliable estimation. When few observations are available, SEs will be large and inaccurate, leading to low statistical power. Further, resulting interval estimates could span values less than zero or greater than one, or have zero width which is practically meaningless. Second, when mislabelling rates are exactly 0% or 100%, SEs will be exactly 0%. For species like snapper, where 100% mislabelling rates are often found, the above equation is completely useless. In terms of sample collection, market survey designs currently lack statistical rigor, which generally manifests as convenience sampling along with pseudoreplication. For example, many fish fillets purchased in supermarkets can be traced back to a single supplier. Therefore, more sophisticated approaches are warranted.

A controversial study by Stawitz et al. (2016) that examined both the financial impact and the ecological ramifications of global seafood product fraud for a variety of fish species (6754 samples spanning 43 DNA barcoding publications) employed a frequentist Generalised Linear Model (GLM) framework in combination with model selection using the Akaike Information Criterion (AIC) (Akaike, 1974) given by

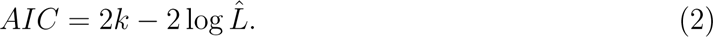

where *k* is the number of parameters and *L*^ is the MLE. The authors utilised taxonomic information, in addition to species price data and species conservation status to develop logistic regression models in an effort to determine the overall effects of seafood mislabelling for samples collected from six sources: port, distributor, grocery, market, restaurant, and sushi (Stawitz et al., 2016). However, there were several major limitations to this work. The best GLM in terms of lowest AIC reported by Stawitz et al. (2016), which incorporated genus, country and purchase source information as covariates for the five most commonly consumed finfish groups in the USA (Atlantic and Pacific salmon, tuna, catfish, and cod), found that species tested using DNA barcoding were both cheaper (-2.98% ex-vessel price) and of higher conservation status (+3.88% IUCN conservation status) compared to the species names listed on product labels. Firstly, the use of the IUCN Red List status was very crude. For products that could be identified to the genus or species level, which accounted for 32.520% of samples, the authors used the average status of the respective species, which is entirely inaccurate, due to varying conservation status at the genus level. This led the researchers to conclude that a label can often be of more severe conservation status than the true species, which was contradictory to most other research. The authors argued that if species of lower conservation priority are being used to substitute species of higher conservation concern, then product mislabelling might actually be increasing seafood sustainability. From an interpretive perspective, in reality, this is far from the truth. Stawitz et al.’s (2016) findings and overall conclusions were immediately called into question in a number of rebuttals (Donlan et al., 2017; Mariani et al., 2017; Warner et al., 2017). Criticism revolved around the application of the USFDA Seafood List to inform global species mislabelling trends and the use of inappropriate GLM covariates to estimate mislabelling rates, among other misapplications. In response to unfavourable attention generated from their earlier work, especially by Donlan et al. (2017), who noted “seafood fraud deserves better”, Stawitz et al. (2016) admit clear statistical oversights.

Other studies have employed statistical regression methods in an effort to better assess global seafood mislabelling impacts at many hierarchical levels, but most are either inconsistent in methodology, or incomplete in analysis. Most studies simply employ basic one-way or two-way tests of independence or goodness of fit, but several studies have implemented more advanced methods. Kim and Lee (2018) utilised ordered probit regression to model consumer purchasing and consumption preference to eco-labelled seafood (flatfish, salmon, tuna, and octopus) across 2773 households in Korea, accounting for covariates such as seafood freshness and price, as well as age, sex, and income of survey respondents. Munguia-Vega et al. (2022) used log-transformed linear models to investigate the global and local influences on patterns of mislabelling and substitution in the Mexican fish trade sector. Perhaps to amend Stawitz et al.’s (2016) misunderstandings, Luque and Donlan (2019) conducted a large-scale Bayesian meta-analysis of global seafood fraud studies totalling 27,313 samples across 141 usable papers (both peer-reviewed and unrefereed) at seven levels including country, source (restaurant, port, market, and grocery), year, product form (processed and fillet), and taxonomic identity (genus and species). Using a two-stage hierarchical modelling framework, an overall product-level mislabelling rate of 8% (95% HPDI (Highest Posterior Density Interval): 4-14%) was noted.

In an effort to place seafood fraud on more statistical footing, here, GLMs within the frequentist and Bayesian inference paradigms are employed to develop a model using predictors known to be informative in the detection of fraud within the Canadian supply chain. A case study conducted by Hu et al. (2018) on mislabelling of finfish products in Metro Vancouver is employed as an illustration to show the overall utility of a rigorous statistical framework for mislabelling estimation, prediction, and classification.

## 2 How can a statistical modelling framework of seafood fraud be employed?

Frequentist and Bayesian statistical inference can be utilised to fulfill numerous short-term and long-term goals of importance to interested parties in several ways, in particular to:

- Capture geographic variability of seafood species fraud (*e.g.*, adulteration, mislabelling, substitution *etc.*) within the supply chain for metropolitan cities across Canada and the USA for the most problematic species, which include cod, halibut, salmon, sole, snapper and tuna;
- Review and evaluate existing trends in seafood product fraud for widely consumed species in terms of biomass availability, market price, and conservation status;
- Employ and adopt developed models in academic, regulatory and collaborator settings to project seafood fraud time of occurrence within major and minor Canadian and US cities for widely misrepresented species at different levels of the supply chain;
- Extend consideration to other geographic regions besides Canada and the USA, particularly those in coastal areas of Europe and Asia, to name a few;
- Develop improved supply chain traceability detection measures and control strategies/guidelines leading to lower prevalence and incidence of seafood fraud in the Canadian and American markets and elsewhere;
- Implement robust policy and legislation to combat seafood fraud based on statistically defensible model predictions of mislabelling rates and time of occurrence for species of economic, environmental, and health concern.

## 3 Case Study: Uncovering Seafood Mislabelling Trends in Metro Vancouver, British Columbia, Canada

To illustrate our arguments above, previously published work by Hu et al. (2018) investigating seafood mislabelling in Metro Vancouver, British Columbia, Canada serves as a case study.

### 3.1 The Dataset

A total of 285 finfish samples was collected randomly or based on product availability from various grocery stores (G; *n* = 93), restaurants (R; *n* = 84), and sushi bars (S; *n* = 108) across Metro Vancouver (comprising Vancouver, Burnaby, Coquitlam, Surrey, Langley and North Vancouver, among others) from September 2017 to February 2018. For samples purchased as takeout from restaurants and sushi bars, taxonomic identity was verified through consultation with establishment employees. Samples were subjected to Polymerase Chain Reaction (PCR) amplification and DNA sequencing using standard laboratory protocols (Hu et al., 2018). Obtained DNA barcode sequences (Hebert et al., 2003a,b) (272 full length 652 basepair (bp) barcodes and nine minibarcodes) were then matched to known references found within the Barcode of Life Data Systems (BOLD; http://www.boldsystems.org) (Ratnasingham and Hebert, 2007). Specimen identifications of product samples gleaned from DNA barcoding were taken at face value even though marker length variation is expected to affect taxonomic resolution depending on the primers utilised for sequence amplification, or the level of sample processing applied after catch (*e.g.*, cooking/canning). For instance, in Wong and Hanner (2008), species mislabelling was based on a DNA sequence similarity top match threshold of 97%. That is, barcodes were deemed to originate from different species whenever sequences found in seafood markets differed by at least 3% from those in BOLD or GenBank (https://www.ncbi.nlm.nih.gov/genbank/). Because labelling status was missing for four samples (G80, R2, R51, and S10), as they failed to yield either a full length or mini DNA barcode sequence (Hu et al., 2018), these data were excluded from the model and instead utilised for downstream prediction and classification (**Table 1**); see section **3.2.2** for further details.

**Table 1:** Characteristics of seafood samples with unknown labelling status.

| Sample | State | Appearance | Form | Colour |
| --- | --- | --- | --- | --- |
| G80 | Raw | Modified | Fillet | Light |
| R2 | Cooked | Modified | Fillet | Red |
| R51 | Cooked | Modified | Fillet | Light |
| S10 | Raw | Plain | Fillet | Light |

While data exclusion is often frowned upon, since it can lead to a loss of statistical power to detect real effects and potential estimation bias, missing data accounted for only 1.404% of the dataset. Thus, 281 samples (representing 98.6% of the entire dataset) were employed for all subsequent analyses (see **Supplementary Tables 1-5** for a breakdown by labelling status (*i.e.*, correctly labelled or mislabelled)). Collected samples were assessed based on the following characteristics: Source (grocery (*n* = 92), restaurant (*n* = 82), and sushi bar (*n* = 107)), State (cooked (*n* = 87) and raw (*n* = 194)), Appearance (modified (seasoned, or covered with breading/other ingredients; *n* = 124) or plain fillet (*n* = 157)), product Form (chopped (*n* = 3), chunk (*n* = 27), fillet (*n* = 244) and whole (*n* = 7)) and Colour of the flesh (light (*i.e.*, pink/white; *n* = 177) or dark (*i.e.*, red; *n* = 104)). Findings by Hu et al. (2018) revealed an overall average mislabelling rate of *p*^ = 25% (70/281; *S*^’^*E*[*p*^] = 0.026 by Equation (1); 29% in restaurants, 24% in grocery stores, and 23% in sushi bars). Three species possessed the highest mislabelling rates: snapper (31/34; 91.170%), halibut (6/24; 25%), and sole (3/19; 15.790%) (Hu et al., 2018). In addition, Hu et al. (2018) implemented a binary logistic GLM in MATLAB (The MathWorks Inc., 2022) using all surveyed covariates as predictors to assess mislabelling of collected seafood products. Obtained significance test results can be found in Table 3 of their study.

### 3.2 The Model, Analysis, and Overall Findings

Hu et al.’s (2018) study leaves much to be desired on statistical grounds as many questions surrounding seafood mislabelling practices currently remain unanswered. In an effort to verify results of their GLM and validate claims made by the authors, data were refitted herein. As with Hu et al. (2018), model interactions are not considered here due to increased model complexity and interpretability issues, despite strong interactions among predictors being conceivable. For instance, a positive Source *×* Appearance interaction term which is statistically significant could suggest that the effect of Source on seafood mislabelling rates is stronger when the appearance of the seafood is altered. This in turn could suggest that seafood samples obtained from restaurants may be more likely to be mislabelled when their appearance is modified compared to when it is plain. Thus, the incorporation of interaction effects is left for future work. The single-intercept (*β*_0_), no-interaction logistic regression (logit) model employed herein is given by:

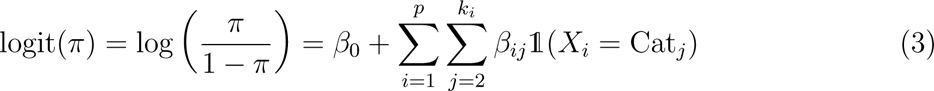

where *π*, the success probability (here, the seafood mislabelling rate), forms the basis of the remainder of the paper. The quantity *<u>^π^</u>* is the odds in favour of product mislabelling relative to a sample being correctly labelled, *p* is the total number of predictors, *X_i_* represents the *i*th categorical predictor, *k_i_*is the number of categories (levels) for the *i*th predictor, *β_ij_* represents the coefficient for the *j*th category of the *i*th predictor, and `1`(*X_i_* = Cat*_j_*) is an indicator function that takes the value one when *X_i_* is in the *j*th category, and zero otherwise. Probabilities are then computed from a simple transformation:

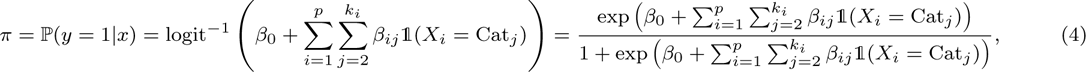

which makes results more easily interpretable. Specifically, the model comprised fixed effects for both intercept and regression coefficients; however, each factor level associated with a given factor (*e.g.*, grocery, restaurant, and sushi bar in the case of Source) will not always have the same effect. Within Equations (3) and (4), a total of *k* = 9 parameters must be estimated (including the intercept and excluding baseline levels for each factor; see below), with *p* = 5 representing the source, state, appearance, form, and colour of retrieved seafood samples. Thus, model parameters must be estimated numerically either through frequentist methods via Iteratively Reweighted Least Squares (IRLS) to obtain MLEs, or through Bayesian inference via the posterior distribution, each based on available data. Unlike in most standard regression scenarios, here the intercept is directly interpretable since all covariates are categorical: *β*_0_ is the log odds of mislabelling when all predictors are at their reference level. By definition, reference levels correspond to *β_i_* = 0, giving an odds of 1 and a probability of 50%. In the context of the present case study, *β*_0_ is the estimated mislabelling log odds associated with a cooked modified chopped light-coloured seafood sample collected from a grocery store. Logistic regression coefficients should be interpreted in the following way: for every one unit increase in a given predictor variable *x_i_*, the log odds increases additively by *β_i_* units, given all other predictor variables are held constant. This, in turn, corresponds to a change in the odds by a multiplicative factor of exp(*β_i_*) units. While logistic regression coefficients are not straightforward to decipher, especially in the presence of categorical variables and interactions, it is important not to associate estimated effects with true causation. Herein, in addition to estimation, probabilities obtained from model fitting are used to classify the four samples having unknown labelling status as either mislabelled (1) or correctly labelled (0) according to

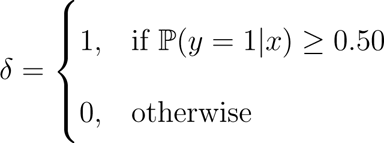

That is, samples with predicted probabilities exceeding 50% are likely to be mislabelled compared to those below this threshold.

All analyses in the present work were carried out on a 2023 Apple MacBook Pro with M2 chip and 16 GB RAM running macOS Ventura 13.2. A random seed was set to guarantee reprodicibility of obtained results. Outputted regression estimates were rounded to three decimal places of precision and interpreted as absolute percentage differences from baseline values of 0 and 1 for log odds and odds, respectively, as well as 50% for probabilities.

#### 3.2.1 Frequentist Analysis

Since Hu et al. (2018) did not report regression estimates nor derived statistics thereof, their model was re-examined in R via glm() using the binomial family with the default logit link function, outputting log odds, odds, and probabilities, among other important quantities contained in the summary() function (**Tables 2** and **3**). In addition to these, associated asymmetric 95% profile likelihood CIs were also computed using the confint() function, along with monotonic transformation due to the problems inherent with symmetric Wald intervals calculated via Equation (1) discussed previously. Reference levels for each factor included in the model were coded alphabetically (which is the default in R). For instance, “Grocery” served as the reference for Source since it occurs before both “Restaurant” and “Sushi bar”. In the case of *L* levels, there are *L −* 1 dummy/indicator variables used to represent the categories of a factor. Typically, reference levels are designated based on the most representative factor level, or that which is most important to a given research question. However, because so little is known about the extent of seafood fraud within the supply chain, custom coding of reference levels was not utilised. This was followed by testing for multicollinearity among predictors via the vif() function in the car package (Fox and Weisberg, 2019) using default argument settings. vif() computes Variance Inflation Factors (VIFs) based on regressing each predictor onto all others and then reporting the coefficient of determination (*R*^2^). The VIF is given by:

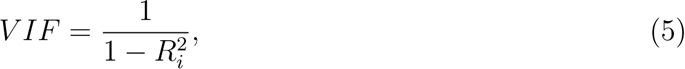

where variables with VIF *>* 5 (equivalent to an *R*^2^ *>* 0.800) should be dropped from the model. When regression models contain many predictors, there is a strong chance that a number of covariates are perfectly correlated (having correlation 1 or -1) with one another. This can lead to identifiability issues since one predictor is a linear function of the other. While some main effects were found to not be significant at the 5% level (**Table 2**), such predictors were not removed from the model due to their perceived importance in the seafood fraud detection literature and the desire to replicate past findings. This said, it would be worthwhile as part of future work to employ model selection procedures, such as stepwise regression using criteria like the AIC, to arrive at a simpler and more explanatory model. Multicollinearity was dealt with through calculation of the generalised variance inflation factor (GVIF) which is an extension of the VIF for models containing categorical variables. Here, a scaled version of the GVIF, sGVIF, is employed, which is given by:

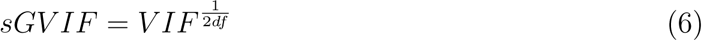

**Table 2:**
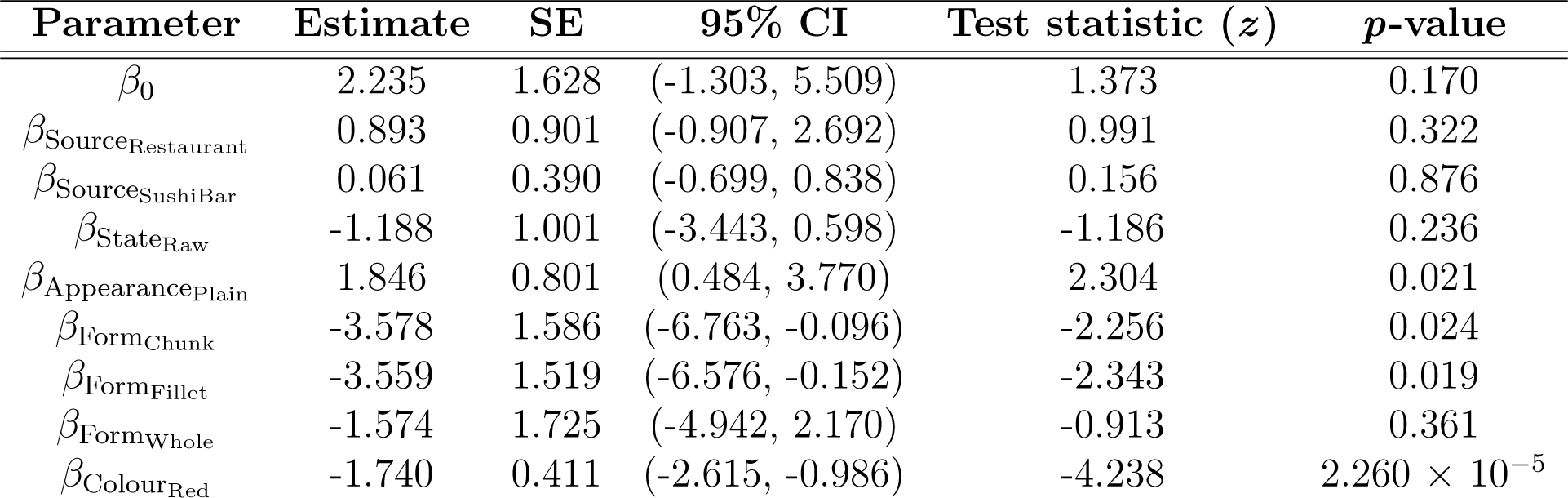
Frequentist GLM summary results and CIs computed via Maximum Likelihood. The Estimate column denotes the log odds relative to the reference level, whereby values are compared to a baseline of zero.

| Parameter | Estimate | SE | 95% CI | Test statistic ($z$) | $p$-value |
| --- | --- | --- | --- | --- | --- |
| $\beta_0$ | 2.235 | 1.628 | (-1.303, 5.509) | 1.373 | 0.170 |
| $\beta_{\text{SourceRestaurant}}$ | 0.893 | 0.901 | (-0.907, 2.692) | 0.991 | 0.322 |
| $\beta_{\text{SourceSushiBar}}$ | 0.061 | 0.390 | (-0.699, 0.838) | 0.156 | 0.876 |
| $\beta_{\text{StateRaw}}$ | -1.188 | 1.001 | (-3.443, 0.598) | -1.186 | 0.236 |
| $\beta_{\text{AppearancePlain}}$ | 1.846 | 0.801 | (0.484, 3.770) | 2.304 | 0.021 |
| $\beta_{\text{FormChunk}}$ | -3.578 | 1.586 | (-6.763, -0.096) | -2.256 | 0.024 |
| $\beta_{\text{FormFillet}}$ | -3.559 | 1.519 | (-6.576, -0.152) | -2.343 | 0.019 |
| $\beta_{\text{FormWhole}}$ | -1.574 | 1.725 | (-4.942, 2.170) | -0.913 | 0.361 |
| $\beta_{\text{ColourRed}}$ | -1.740 | 0.411 | (-2.615, -0.986) | -4.238 | $2.260 \times 10^{-5}$ |

**Table 3:** Frequentist GLM predictor odds, probabilities, and CIs derived via the Estimates column in Table 2. Odds above one indicate an increase in overall sample mislabelling, whereas odds below one signify a decrease relative to the baseline. A similar interpretation exists for probabilities at a predefined decision threshold of 50%.

| Parameter | Odds | 95% CI | Probability | 95% CI |
| --- | --- | --- | --- | --- |
| $\beta_0$ | 9.346 | (0.272, 246.903) | 0.903 | (0.214, 0.996) |
| $\beta_{\text{SourceRestaurant}}$ | 2.443 | (0.404, 14.766) | 0.710 | (0.288, 0.937) |
| $\beta_{\text{SourceSushiBar}}$ | 1.063 | (0.497, 2.312) | 0.515 | (0.332, 0.698) |
| $\beta_{\text{StateRaw}}$ | 0.305 | (0.032, 1.818) | 0.234 | (0.031, 0.645) |
| $\beta_{\text{AppearancePlain}}$ | 6.332 | (1.623, 43.378) | 0.864 | (0.619, 0.977) |
| $\beta_{\text{FormChunk}}$ | 0.028 | (0.001, 0.909) | 0.027 | (0.001, 0.476) |
| $\beta_{\text{FormFillet}}$ | 0.028 | (0.001, 0.859) | 0.027 | (0.001, 0.462) |
| $\beta_{\text{FormWhole}}$ | 0.207 | (0.007, 8.755) | 0.171 | (0.007, 0.897) |
| $\beta_{\text{ColourRed}}$ | 0.175 | (0.073, 0.373) | 0.149 | (0.068, 0.272) |

where *df* is the number of coefficients in a subset of the full model. Given sGVIFs were all less than five, each predictor was kept in the model (**Supplementary Table 6**). Despite this, sGVIFs for both Appearance and Form were larger than those for the remaining covariates, suggesting that perhaps one of these predictors is problematic and therefore should be excluded. This is left for a followup study. A classic problem encountered when performing logistic regression with categorical variables in a frequentist setting is separation, which occurs when a linear combination of the data classes (here, *e.g.*, mislabelled and correctly labelled) are perfectly predicted by underlying models (Mansournia et al., 2017). In the context of seafood mislabelling, this can occur, for example, whenever all samples belonging to a particular species have a 0% mislabelling (similarly, correctly labelled) rate or a 100% mislabelling (correspondingly, correctly labelled) rate, or when sample sizes for predictor categories are low. When this happens, parameter estimates and/or SEs will be very large or infinite due to lack of algorithm convergence. A common fix to rectify non-identifiability in this regard is to prune predictors until the model is identifiable; however, this could result in the exclusion of highly predictive variables from the final model. Such an issue was not encountered within the study herein, as no warnings were returned by glm() during fitting, and all computed coefficient estimates and SEs exhibited numeric stability. Fortunately, the Bayesian paradigm offers a simple way forward without the need to collect additional data through techniques like model/parameter hierarchicalisation, where specification of suitable priors can effectively regularise coefficients by shrinking them toward zero.

In comparing frequentist GLM results obtained herein to Hu et al. (2018), several findings are noteworthy. Here, only coefficients for plain appearance (*β*_AppearancePlain_ ), chunk form (*β*_FormChunk_ ), fillet form (*β*_FormFillet_ ), and red colour (*β*_ColourRed_ ) were found to be statistically significant (*p ≤* 0.05) (**Table 2**), whereas *β*_0_, *β*_form_, and *β*_colour_ were in the case of Hu et al. (2018). Unfortunately, since Hu et al. (2018) did not provide specific modelling details (only *F* -test results were given), nor coding scripts, exact reproducibility of their findings is challenging. It is also important to note that unlike R, which is free and open-source software, MATLAB is proprietary and subscription-based. In particular, it is unclear what factor levels were employed by Hu et al. (2018) as references. However, like R, MATLAB uses the first level listed alphabetically to denote the reference category; thus, it is assumed that this was the case for Hu et al. (2018). Regardless, statements made by Hu et al. (2018) concerning their results are too general to be practicably applicable, and also lack strong statistical support. Hu et al. (2018) provide no justification for the reporting of *F* -test *p*-values. In the context of logistic regression, a deviance test is commonly conducted to assess model fit; yet Hu et al. (2018) did not carry out any regression model comparison. Phrasing in Hu et al. (2018) at times is also ambiguous and inconsistent, as it was unclear how comparisons were being made: “Products composed of fish muscle of a lighter color flesh and chopped muscle tissue were more prone to illicit practices compared to red or fillet samples…”. Furthermore, the use of catchall terms like “source” and “appearance” in Table 3 of Hu et al. (2018) strongly suggests that the authors coded their model improperly and likely treated categorical variables as integer values. Had the authors likely not made this error, compared to results within the present study, theirs would still differ due to the fact that default optimisation algorithms, parameter settings, numerical precision, and random seeds used for estimation, among other factors, all differ between R and MATLAB. Findings herein are revealing: a cooked modified chopped light coloured seafood sample collected from a grocery store corresponds to a 223.492% higher log odds, 834.574% increased odds and 40.334% greater chance of being mislabelled at a threshold of 50% compared to non-reference samples.

Further, product source (*β*_SourceRestaurant_ and *β*_SourceSushiBar_ ) is associated with higher log odds, odds and probabilities of mislabelling relative to grocery store samples (89.308%, 144.264%, and 20.953%, and 6.103%, 6.293%, and 1.525%, respectively) (**Tables 2** and **3**). Similarly, *β*_AppearancePlain_ has a 184.559% higher log odds, 533.182% increased odds, and 36.361% greater probability at a 50% boundary of being mislabelled compared to modified products. Note that since summary() carries out a two-sided significance test at level *α* (herein set at 5%) by default, there is a direct relationship between hypothesis tests and reported 95% CIs: if a hypothesised value of a population parameter (herein zero, one, and 50% for log odds, odds, and probabilities, respectively) falls within said interval, then the null hypothesis of equality will fail to be rejected. Hu et al. (2018) point to sampling bias as a likely culprit, leading to artificially high mislabelling rates due to low numbers of collected samples. This can be seen in the Form category in the case of chopped, chunk, and whole samples (**Supplementary Table 4**). Conversely, *β*_StateRaw_ , all Form parameters, and *β*_ColourRed_ each have a 118.759%, 69.505% and 26.632%, 357.802%, 97.207%, and 47.283%, 355.850%, 97.152%, and 47.231%, 157.447%, 79.288%, and 32.842%, and 174.047%, 82.456%, and 35.075% lower log odds, odds and probabilities of mislabelling relative to cooked, chopped, and light coloured samples, respectively.

#### 3.2.2 Bayesian Analysis

The model was then fitted using the Stan probabilistic programming language (Carpenter et al., 2017) framework with default settings for leapfrog integration (*e.g.*, step size) via the No-U-Turn Sampler (NUTS) algorithm (Hoffman and Gelman, 2014). The rstanarm R package (Goodrich et al., 2023) was employed to avoid needing to first compile code in the rstan R package (Stan Development Team, 2023a), to ensure generation of well-behaved Markov chains, and to aid ease of reproducibility. Unlike frequentist inference, its Bayesian counterpart treats unknown parameters (*β*) as random variables proportionally through Bayes’ theorem to derive a joint posterior distribution from the likelihood (*p*(*y|X, β*)) and an *a priori* chosen prior distribution (*p*(*β*)):

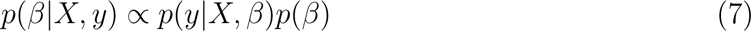

(Gelman et al., 2014; van de Schoot et al., 2021), where equality holds through division of the right-hand side of the above expression by the prior predictive distribution *p*(*y|x*) = *p*(*y|X, β*)*p*(*β*) d*β*, in which model parameters are marginalised out. However, often this integral has no analytical (*i.e.*, closed-form) solution, thus necessitating numerical approximations. Therefore, Hamiltonian Monte Carlo (HMC) was employed to generate dependent samples from the posterior distribution of interest. HMC is inspired by Hamiltonian mechanics, wherein a proposed sample moving toward the target distribution is likened to the movement of a particle in a physical system based on its position, velocity, energy, and momentum over time (Neal, 2011). Compared to other Markov Chain Monte Carlo (MCMC) algorithms like the Metropolis-Hastings sampler (Hastings, 1970) and the Gibbs sampler (Geman and Geman, 1984), HMC sampling in Stan is much faster computationally and is much more efficient for challenging posteriors in terms of chain mixing compared to sister software like JAGS (Just Another Gibbs Sampler) (Plummer, 2003). This is the case since HMC relies on gradient (*i.e.*, multivariate derivative) information to explore the parameter search space, whereas JAGS imposes no such overhead, as it is essentially a random walk. Further, because resulting samples are minimally autocorrelated, Markov chain thinning is not required. Thus, convergence to target densities is quite rapid and acceptance rates of proposed samples is optimal. This behaviour is especially desirable in high-dimensional situations, where the number of model parameters exceeds the number of sampled observations. In the context of seafood fraud, product mislabelling rate estimation can be viewed as a high-dimensional problem as few samples are often collected across multiple predictors (*e.g.*, source, geographical location, time).

A variety of prior distributions on model parameters available in rstanarm was examined here to assess model sensitivity: (1) a *N* (0, 2.5) prior on the intercept with non-informative (flat) uniform *U* (*−∞, ∞*) priors on regression coefficients, (2) weakly informative standard Normal *N* (0, 1) priors on all model parameters, (3) weakly informative Normal *N* (0, 2.5) priors on all model parameters and (4) weakly informative Cauchy priors on all model parameters. Prior (1) above is specified in the absence of any information regarding the true values of posterior parameters, whereas prior (2) is a common choice widely seen in the literature. Prior (3) is rstanarm’s default, which tends to work well in most scenarios. The choice of prior (4) requires some explanation. The Cauchy distribution has been recommended in the context of logistic regression by several authors (Gelman et al., 2014, 2008; Ghosh et al., 2018), as well as in the Stan User Guide (Stan Development Team, 2023b) and by the Stan Development Team (Stan Development Team, 2020), in part due to the distribution’s heavier tails compared to the Normal distribution, allowing the incorporation of outliers. In particular, Gelman et al. (2008) remarks that logistic regression coefficients rarely lie outside the interval (-5, 5), which corresponds to approximately (0.007, 0.993) on the probability scale by Equation (4). As such, a *Cau*(0, 10) distribution is recommended on the model intercept and a *Cau*(0, 2.5) on all other coefficients (Gelman et al., 2008). For all considered priors, model intercepts coincide with predictors centered to have mean zero. Centering in this fashion can often facilitate interpretation of models in the presence of interaction effects (Gelman, 2007).

Four chains were run for 2000 iterations each in parallel across four cores. Within each chain, a total of 1000 samples was discarded as warmup (*i.e.*, burnin) to reduce dependence on starting conditions. Further, 1000 post-warmup draws were utilised per chain. Each of these reflect default MCMC settings in Stan. MLEs for regression coefficients given in **Table 1** were used as initial values for MCMC. Many past studies employing Bayesian analysis centre prior distributions on MLEs; however, this “double dipping” is bad practice for two reasons (van de Schoot et al., 2021). First, priors should be specified independently of the data (*i.e.*, before seeing any data or a statistical summary thereof). Secondly, centering of prior distributions results in inflation of posterior effective sample sizes (ESSs; see below) relative to actual parameter sample sizes, leading to overconfidence in obtained results. Note, Stan parameterises models using the standard deviation, not the variance. Convergence was assessed in a number of ways as suggested by Gelman et al. (2020): (1) through examining parameter traceplots, which depict the trajectory of accepted MCMC draws as a function of the number of iterations, (2) through monitoring the Gelman-Rubin *R*^^^ statistic (Gelman and Rubin, 1992; Vehtari et al., 2021), which measures within-chain *versus* between-chain variance, and (3) through calculating the ESS for each parameter, which quantifies the number of independent samples generated Markov chains are equivalent to. Mixing of chains was deemed sufficient when traceplots looked like “fuzzy caterpillars”, *R*^^^ *<* 1.01, and effective sample sizes were reasonably large. Alongside sample posterior means and posterior standard deviations (SDs), 95% credible intervals (CrIs) were also reported, which should be interpreted as follows, to the extent that one trusts the prior distribution selected and underlying data: Given a (1 *− α*)100% CrI, where *α* is the desired significance level, the true parameter is contained within said interval with (1 *− α*)100% probability. This is in sharp contrast to a CI, where, with repeated sampling, on average (1 *− α*)100% of constructed intervals will contain the true parameter of interest.

Selected priors were deemed appropriate based on prior predictive checks generated in rstanarm and the bayesplot R package (Gabry et al., 2019; Gabry and Mahr, 2022) to ensure simulated sample mean draws were reasonably close to observed data; the same was done for resulting posteriors (**Supplementary** Figures 1-8). Results in **Table 4** and **Table 5** highlight that the Bayesian model is sensitive to the choice of prior distribution, but some parameters are affected more than others likely owing to a small number of observations for particular factor levels like those for Form. Interpretation of results in terms of percent deviation from baseline values is identical to that in section **3.2.1**. Further, for all assessed prior distributions, estimates appear close in value to their frequentist counterparts. This is especially true in the case of the uniform prior, since the posterior is completely dominated by the likelihood. Across all examined priors, parameter traceplots showed rapid convergence to the stationary distribution (**Supplementary** Figures 9-12), and effective sample sizes (not shown) were all above 1000. Estimates for many parameters (*e.g., β*_0_) are consistent yet highly uncertain across priors, given large posterior SDs and wide CrIs. Despite this, CIs and CrIs overlap considerably across priors. This indicates that the seafood mislabelling problem is not as simple as it appears to be on the outset for Bayesian computation to solve. Such a finding strongly suggests that prior elicitation from a domain-specific expert is likely warranted and would be beneficial, as no clear prior seems to outperform any other across all estimated model parameters for the data at hand. A path forward could be to employ priors specified by Donlan et al. (2017). Additionally, it stands to reason that expanding the model to the pooled, unpooled and hierarchical cases, as well as those incorporating spatiotemporal structure, each with fixed/varying intercepts and varying regression coefficients, would likely lead to better agreement across selected priors, as well as more realistic interpretation. In the case of a hierarchical model, a common distribution for hyperparameters and hyperpriors could be easily specified. However, this could result in a lower degree of interpretability due to the adoption of a more complicated model with a larger number of parameters estimated from relatively little data, as well as sampling pathologies commonly encountered in HMC when exploring posterior parameter spaces possessing complex geometries (*e.g.*, curvature). In the latter case, divergent transitions commonly arise, which may prove difficult to diagnose and resolve. A path forward could be to increase the target average acceptance probability (adapt delta), which is set to 0.800 by default in rstan, to a value like 0.999. Doing so will result in smaller step sizes taken by the leapfrog integrator, in addition to increasing computational expense. However, the end result will be more robust due to more efficient sampling. Another approach to remedy HMC sampling difficulties, especially in hierarchical settings, is model reparameterisation, in particular centering, which can often achieve convergence in fewer iterations and with larger effective sample sizes (Betancourt and Girolami, 2013); however, implementation can prove challenging.

**Table 4:** Bayesian GLM estimates. Interpretation is identical to that in Table 2.

| Parameter | Flat | SD | 95% CrI | Normal | SD | 95% CrI | Default | SD | 95% CrI | Cauchy | SD | 95% CrI |
| --- | --- | --- | --- | --- | --- | --- | --- | --- | --- | --- | --- | --- |
| $\beta_0$ | 2.271 | 1.952 | (-1.810, 5.907) | -0.354 | 0.811 | (-1.970, 1.213) | 1.879 | 1.788 | (-1.868, 5.225) | 0.529 | 1.382 | (-2.200, 3.286) |
| $\beta_{\text{SourceRestaurant}}$ | 0.964 | 0.933 | (-0.908, 2.814) | 0.526 | 0.579 | (-0.573, 1.684) | 0.901 | 0.911 | (-0.879, 2.703) | 0.766 | 0.776 | (-0.755, 2.290) |
| $\beta_{\text{SourceSushiBar}}$ | 0.083 | 0.384 | (-0.656, 0.840) | -0.080 | 0.345 | (-0.752, 0.600) | 0.060 | 0.396 | (-0.696, 0.841) | -0.003 | 0.382 | (-0.748, 0.728) |
| $\beta_{\text{StateRaw}}$ | -1.464 | 1.076 | (-3.753, 0.403) | -0.577 | 0.579 | (-1.726, 0.556) | -1.287 | 0.989 | (-3.336, 0.497) | -0.937 | 0.825 | (-2.662, 0.610) |
| $\beta_{\text{AppearancePlain}}$ | 2.171 | 0.920 | (0.597, 4.240) | 0.993 | 0.492 | (0.087, 1.986) | 1.952 | 0.831 | (0.536, 3.790) | 1.513 | 0.694 | (0.285, 3.036) |
| $\beta_{\text{FormChunk}}$ | -3.816 | 1.924 | (-7.493, 0.193) | -0.766 | 0.702 | (-2.198, 0.628) | -3.386 | 1.758 | (-6.703, 0.329) | -1.915 | 1.368 | (-4.739, 0.707) |
| $\beta_{\text{FormFillet}}$ | -3.667 | 1.850 | (-7.101, 0.334) | -0.695 | 0.663 | (-1.961, 0.623) | -3.220 | 1.697 | (-6.423, 0.451) | -1.774 | 1.299 | (-4.333, 0.776) |
| $\beta_{\text{FormWhole}}$ | -1.433 | 2.079 | (-5.236, 3.034) | 0.731 | 0.748 | (-0.733, 2.208) | -1.021 | 1.915 | (-4.630, 3.020) | 0.297 | 1.423 | (-2.383, 3.193) |
| $\beta_{\text{ColourRed}}$ | -1.837 | 0.423 | (-2.718, -1.066) | -1.479 | 0.352 | (-2.196, -0.793) | -1.809 | 0.426 | (-2.696, -1.025) | -1.692 | 0.399 | (-2.505, -0.976) |

**Table 5:** Bayesian GLM odds and probabilities. Interpretation is identical to that in Table 3.

| Parameter | Flat | 95% CrI | Normal | 95% CrI | Default | 95% CrI | Cauchy | 95% CrI |
| --- | --- | --- | --- | --- | --- | --- | --- | --- |
| OddsIntercept | 9.692 | (0.164, 367.652) | 0.702 | (0.139, 3.364) | 6.548 | (0.154, 185.848) | 1.698 | (0.111, 26.746) |
| OddsSourceRestaurant | 2.622 | (0.403, 16.682) | 1.693 | (0.564, 5.385) | 2.463 | (0.415, 14.924) | 2.152 | (0.470, 9.874) |
| OddsSourceSushiBar | 1.086 | (0.519, 2.316) | 0.923 | (0.471, 1.823) | 1.062 | (0.499, 2.319) | 0.997 | (0.473, 2.072) |
| OddsStateRaw | 0.231 | (0.023, 1.496) | 0.562 | (0.178, 1.743) | 0.276 | (0.036, 1.644) | 0.392 | (0.070, 1.841) |
| OddsAppearancePlain | 8.765 | (1.817, 69.424) | 2.698 | (1.091, 7.284) | 7.046 | (1.709, 44.242) | 4.542 | (1.329, 20.830) |
| OddsFormChunk | 0.022 | (0.001, 1.213) | 0.465 | (0.111, 1.873) | 0.034 | (0.001, 1.390) | 0.147 | (0.009, 2.028) |
| OddsFormFillet | 0.026 | (0.001, 1.397) | 0.499 | (0.141, 1.864) | 0.040 | (0.002, 1.569) | 0.170 | (0.013, 2.172) |
| OddsFormWhole | 0.239 | (0.005, 20.774) | 2.078 | (0.481, 9.098) | 0.360 | (0.010, 20.481) | 1.346 | (0.092, 24.354) |
| OddsColourRed | 0.159 | (0.066, 0.344) | 0.228 | (0.111, 0.452) | 0.164 | (0.068, 0.359) | 0.184 | (0.082, 0.377) |
| ProbIntercept | 0.906 | (0.141, 0.997) | 0.412 | (0.122, 0.771) | 0.868 | (0.133, 0.995) | 0.629 | (0.100, 0.964) |
| ProbSourceRestaurant | 0.724 | (0.287, 0.943) | 0.629 | (0.361, 0.843) | 0.711 | (0.293, 0.937) | 0.683 | (0.320, 0.908) |
| ProbSourceSushiBar | 0.521 | (0.342, 0.698) | 0.480 | (0.320, 0.646) | 0.515 | (0.333, 0.699) | 0.499 | (0.321, 0.674) |
| ProbStateRaw | 0.188 | (0.022, 0.599) | 0.360 | (0.151, 0.635) | 0.216 | (0.035, 0.622) | 0.282 | (0.065, 0.648) |
| ProbAppearancePlain | 0.898 | (0.645, 0.986) | 0.730 | (0.522, 0.879) | 0.876 | (0.631, 0.978) | 0.820 | (0.571, 0.954) |
| ProbFormChunk | 0.022 | (0.001, 0.548) | 0.317 | (0.100, 0.652) | 0.033 | (0.001, 0.582) | 0.128 | (0.009, 0.670) |
| ProbFormFillet | 0.025 | (0.001, 0.583) | 0.333 | (0.124, 0.651) | 0.038 | (0.002, 0.611) | 0.145 | (0.013, 0.685) |
| ProbFormWhole | 0.193 | (0.005, 0.954) | 0.675 | (0.325, 0.901) | 0.265 | (0.010, 0.953) | 0.574 | (0.084, 0.961) |
| ProbColourRed | 0.137 | (0.062, 0.256) | 0.186 | (0.100, 0.311) | 0.141 | (0.064, 0.264) | 0.155 | (0.076, 0.274) |

**Table 6:** Predicted frequentist GLM mislabelling log odds, odds, probabilities, and predictions/classifications for missing samples. Labelling status was inferred based on a decision cutoff of 50%. Interpretation of other parameters is identical to that in Tables 2 and 3.

| Sample | log Odds | Odds | Probability | Outcome |
| --- | --- | --- | --- | --- |
| G80 | -2.511 | 0.081 | 0.075 | Correctly labelled |
| R2 | -2.171 | 0.114 | 0.102 | Correctly labelled |
| R51 | -0.431 | 0.650 | 0.394 | Correctly labelled |
| S10 | -0.605 | 0.546 | 0.353 | Correctly labelled |

#### 3.2.3 Prediction and Classification

Estimated mislabelling rates also vary substantially for the four missing samples (**Tables 6** and **7**). The log odds and odds have the same interpretation in terms of percent changes given previously. For instance, in the case of the frequentist GLM, sample R2 (a cooked modified red-coloured fillet collected from a restaurant) is associated with an 88.593% lower odds of being mislabelled compared to a cooked modified chopped light-coloured sample from a grocery store.

**Table 7:**
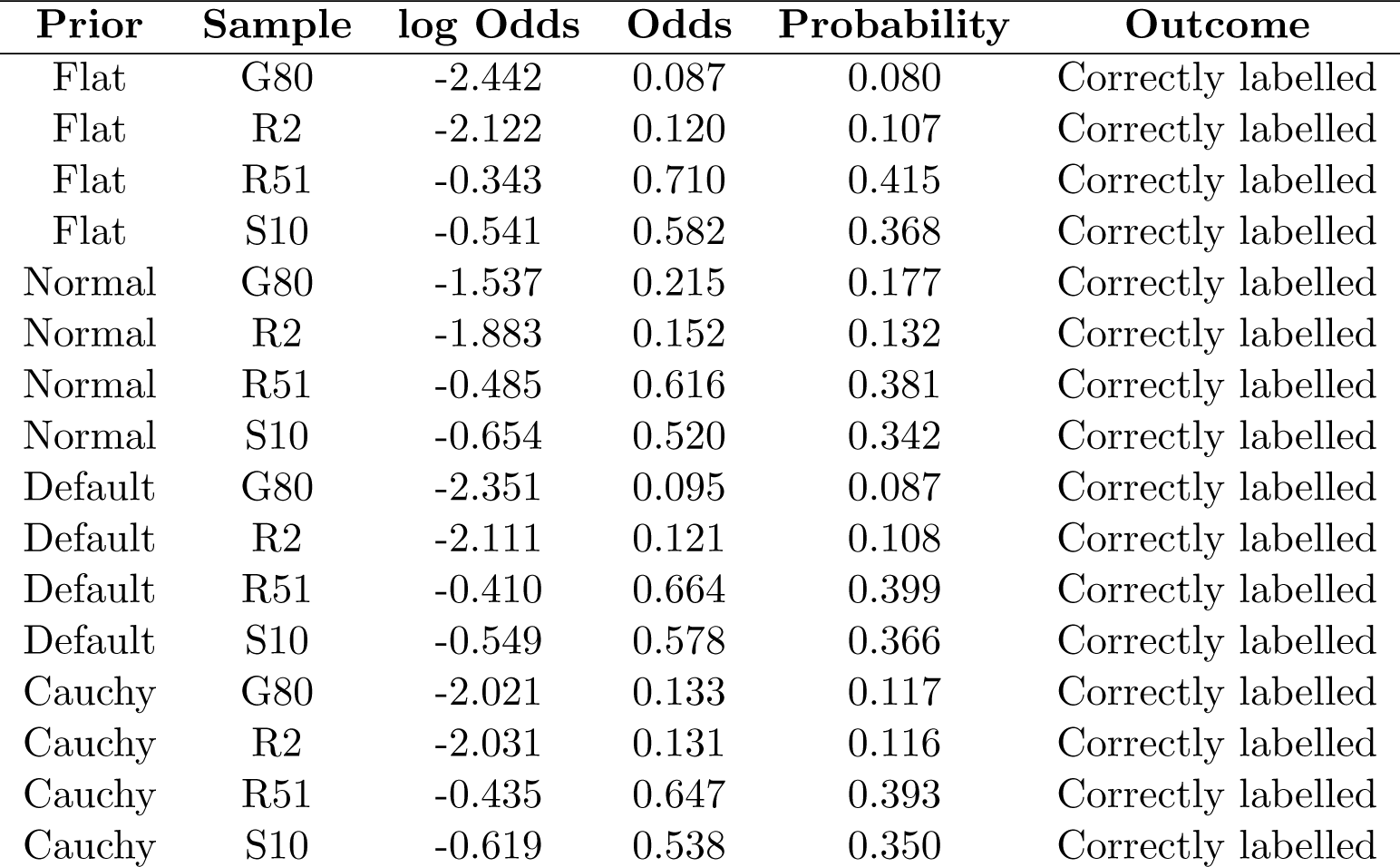
Predicted Bayesian GLM mislabelling log odds, odds, probabilities, and predictions/classifications for missing samples. Labelling status was inferred based on a decision cutoff of 50%. Calculated values are posterior means averaged over 4000 draws. Interpretation of other parameters is identical to that in Table 6.

Overall, the model predicts all missing samples are likely to be correctly labelled under the default decision boundary for both the frequentist and Bayesian GLMs, as all estimated probabilities are below 50%. Despite this, species identification, along with other sample properties, could not be made, as mentioned in section **3.1**. Nevertheless, prediction of samples with unknown labelling status is useful to verify whether findings agree well with observed food processing and fraudulent behaviours, such as removal of morphological features that readily distinguish among species (*e.g.*, skin and scales), concealing at-risk seafood species within highly preserved canned goods, and adding dyes or pigments to farmed Atlantic salmon (*Salmo salar* ) to pass it off as wild-caught Pacific (Sockeye) salmon (*Oncorhynchus nerka*).

As with estimates for model parameters, results depicted in **Tables 6** and **7** demonstrate that the detection of mislabelled seafood is challenging under the chosen priors and default classification threshold, as all estimates are very close in magnitude.

## 4 Conclusion

Herein, a case study on seafood mislabelling trends in Metro Vancouver, British Columbia, Canada by Hu et al. (2018) was utilised to demonstrate the overall promise of frequentist and Bayesian statistical modelling to estimate seafood mislabelling rates, as well as highlight implications for legislation, policy, and health impacts around the import and sale of both common and at-risk species on a global scale. Hu et al. (2018) fell short in their statistical treatment of their data. The fact that their article has been cited nearly 100 times as of January 2024 suggests that findings have not been heavily scrutinized. Herein, it was found that seafood samples obtained from restaurants and sushi bars have a higher chance of being mislabelled compared to those found in grocery stores. Additionally, seafood samples having a plain appearance were over six times more likely to be incorrectly labelled compared to modified samples. Such findings are in line with global trends. For example, product mislabelling rates are often higher in restaurants in comparison to grocery stores because restaurants tend to purchase large volumes of seafood from several different suppliers, distributors, or wholesalers, in sharp contrast to grocery stores who acquire their seafood supply from more homogeneous sources. This in turn leads to problems in transparency, as there are few regulations and limited enforcement specifically targeting restaurants.

While logistic regression was leveraged for the present study, a natural extension to probit regression is entirely feasible. Logit and probit models could then be compared for goodness of fit via the AIC, the Likelihood Ratio Test (LRT), leave-one-out cross validation in a Bayesian context to compare models fitted using a range of prior distributions via the loo R package (Vehtari et al., 2017, 2023), or techniques like Bayesian model averaging/stacking (Yao et al., 2017) to combine plausible models based on their posterior probabilities. Despite this, probit model interpretation is more challenging and less intuitive compared to logistic models, as coefficients are on the *z*-score scale and probabilities must be computed numerically via integration. This is the case since the probit is based on the cumulative distribution function (CDF) of the standard Normal distribution. Furthermore, while the two approaches often give comparable output, estimated probabilities may differ significantly, potentially leading to conflicting prediction outcomes regarding product labelling status. With respect to classification, recall a value of 50% was utilised herein as a decision threshold. Depending on the importance of confidently classifying samples as either correctly labelled or mislabelled (through minimising the number of false positives and false negatives), a different cutoff may be desired. Small thresholds are to be preferred whenever correctly classifying truly mislabelled seafood samples as mislabelled is absolutely critical; however, this comes at a cost of less certainty. Conversely, the use of large cutoffs would result in fewer truly mislabelled samples being detected with greater certainty. To this end, another avenue for future work revolves around classification, where the use of receiver operating characteristic (ROC) curves, which plot the false positive rate (FPR; 1 *−* specificity) against the true positive rate (TPR; sensitivity) and the area bounded by them (AUC), can be utilised on training and testing sets to select optimal thresholds for seafood sample classification found by maximizing Youden’s *J* statistic, given by *J* = sensitivity + specificity *−* 1 (8) in conjunction with measures of performance evaluation such as model accuracy and precision.

Results obtained herein are insightful; however, it is important not to make over-generalisations concerning findings described within the current study: results should only be interpreted in the context of seafood fraud incidence and prevalence within Metro Vancouver. Nevertheless, the integration of frequentist and Bayesian statistical methods into future studies holds much promise.

## Supplementary Information

Information accompanying this article can be found in Supplemental Information.pdf.

## Data Availability Statement

Raw data, R, and Stan code can be found on GitHub at: https://github.com/jphill01/ Phillips-et-al.-Seafood-Fraud-Paper.

## Acknowledgements

We wish to recognise the valuable comments and discussions of Daniel (Dan) Gillis, Robert (Bob) Hanner, and Kathleen (Kat) Nolan along with XXX anonymous reviewers.

We acknowledge that the University of Guelph resides on the ancestral lands of the Attawandaron people and the treaty lands and territory of the Mississaugas of the Credit. We recognize the significance of the Dish with One Spoon Covenant to this land and offer our respect to our Anishinaabe, Haudenosaunee and Métis neighbours as we strive to strengthen our relationships with them.

## Author Contributions

JDP wrote the manuscript, wrote R and Stan code, approved all developed code as well as analysed and interpreted all experimental results. FADVF wrote required R and/or Stan code. All authors contributed to the revision of this manuscript and approved the final version.

