## Supplementary Information for "Statistical modelling of seafood fraud in the Canadian supply chain"

Jarrett D. Phillips and Fynn A. De Vuono-Fraser

### Data Summary

| Factor level | Correctly labelled | Mislabelled |
|---|---|---|
| Grocery (G) | 70 | 22 |
| Restaurant (R) | 58 | 24 |
| SushiBar (S) | 83 | 24 |

**Table 1:** Correctly labelled and mislabelled samples by source.

| Factor level | Correctly labelled | Mislabelled |
|---|---|---|
| Cooked | 62 | 25 |
| Raw | 149 | 45 |

**Table 2:** Correctly labelled and mislabelled samples by state.

| Factor level | Correctly labelled | Mislabelled |
|---|---|---|
| Modified | 96 | 28 |
| Plain | 115 | 42 |

**Table 3:** Correctly labelled and mislabelled samples by appearance.

| Factor level | Correctly labelled | Mislabelled |
|---|---|---|
| Chopped | 2 | 1 |
| Chunk | 24 | 3 |
| Fillet | 183 | 61 |
| Whole | 2 | 5 |

**Table 4:** Correctly labelled and mislabelled samples by form.

| Factor level | Correctly labelled | Mislabelled |
|---|---|---|
| Light | 116 | 61 |
| Red | 95 | 9 |

**Table 5:** Correctly labelled and mislabelled samples by colour.

### Frequentist Analysis

| Variable | GVIF | df | sGVIF |
|---|---|---|---|
| Source | 8.817 | 2 | 1.723 |
| State | 10.290 | 1 | 3.208 |
| Appearance | 6.986 | 1 | 2.643 |
| Form | 1.961 | 3 | 1.119 |
| Colour | 1.094 | 1 | 1.046 |

**Table 6:** Predictor sGVIFs and other related quantities. All predictors have sGVIF $< 5$, indicating little to no evidence of multicollinearity.

### Bayesian Analysis

All results are based on 4000 posterior draws.

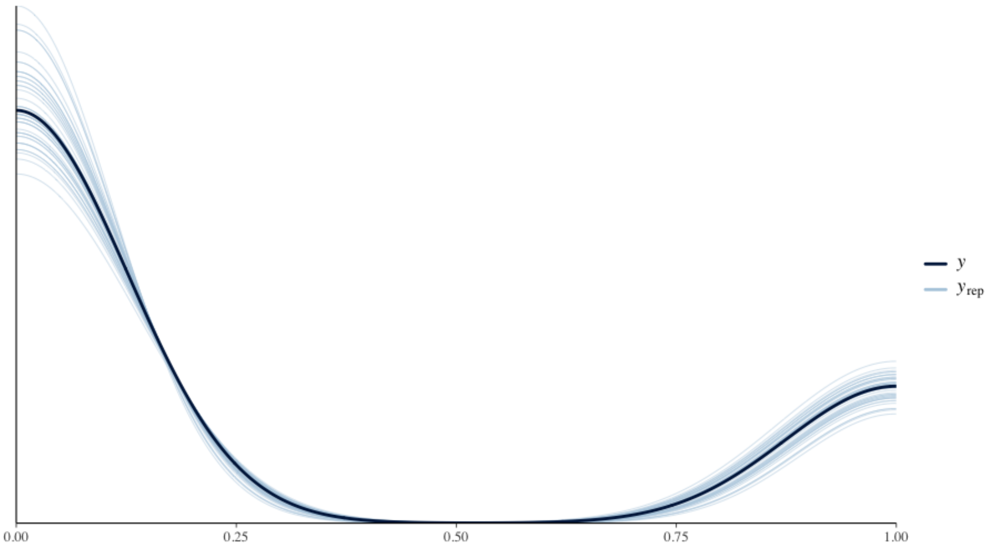


**Figure 1:** Posterior predictive density for $U(-\infty, \infty)$ & $N(0, 2.5)$ prior distribution.

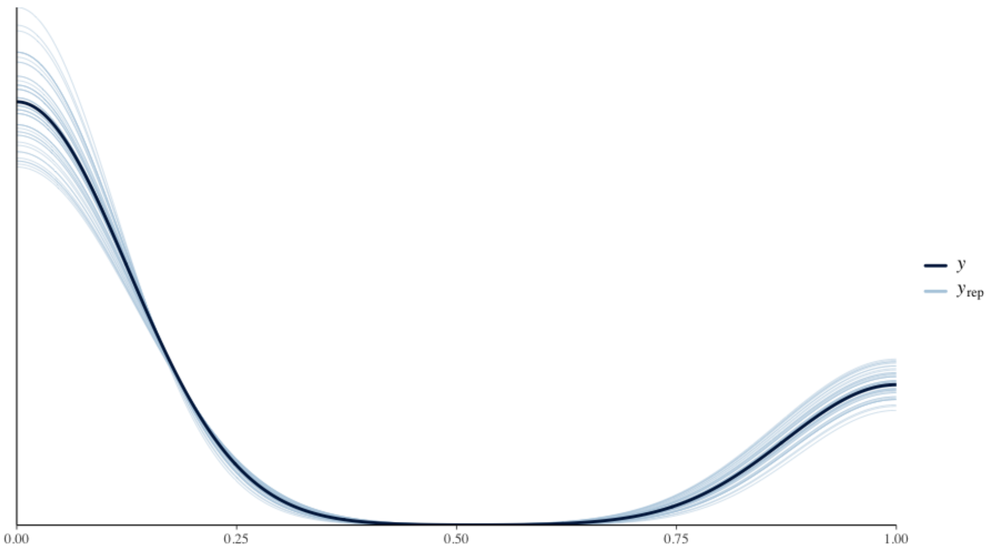


**Figure 2:** Posterior predictive density for $N(0,1)$ prior distribution.

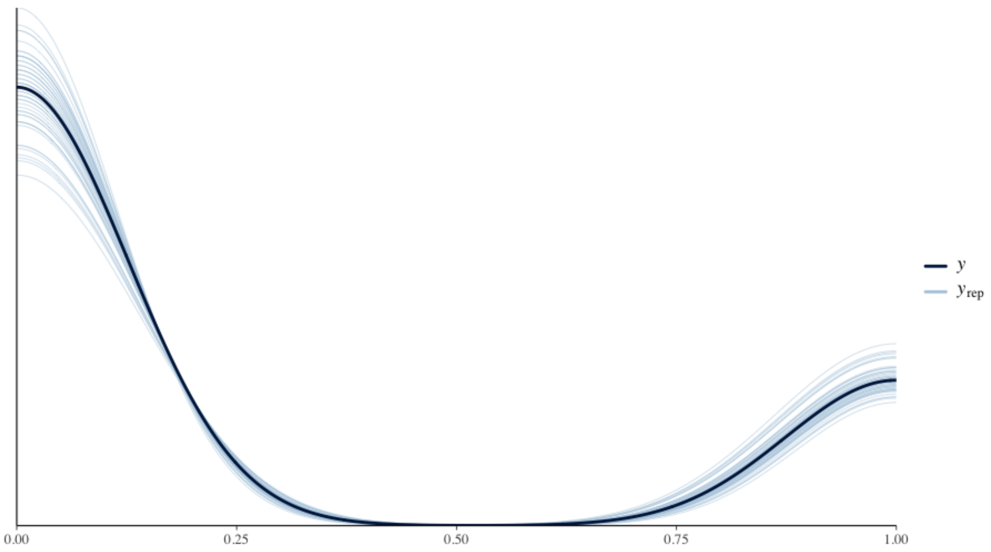


**Figure 3:** Posterior predictive density for $N(0,2.5)$ prior distribution.

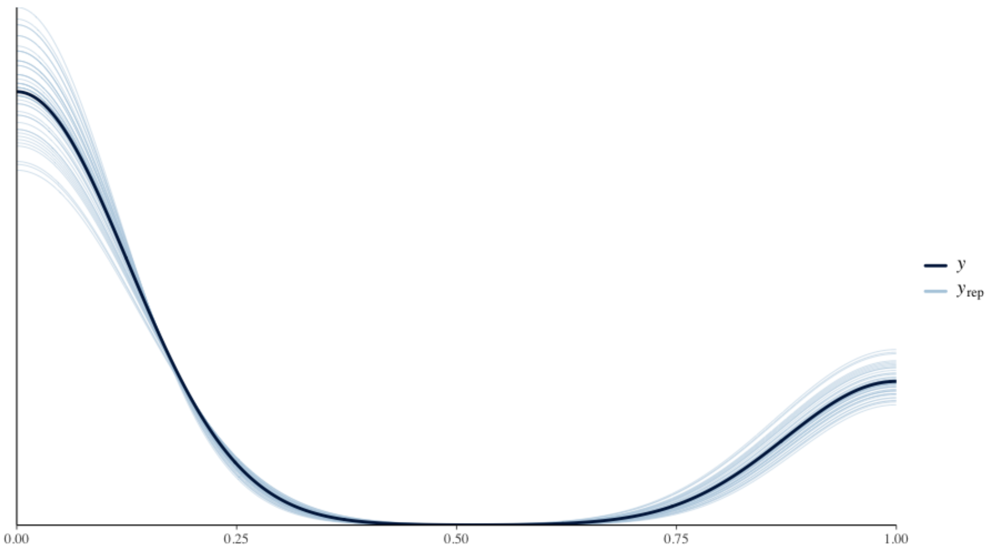


**Figure 4:** Posterior predictive density for $Cau(0, 10)$ & $Cau(0, 2.5)$ prior distribution.

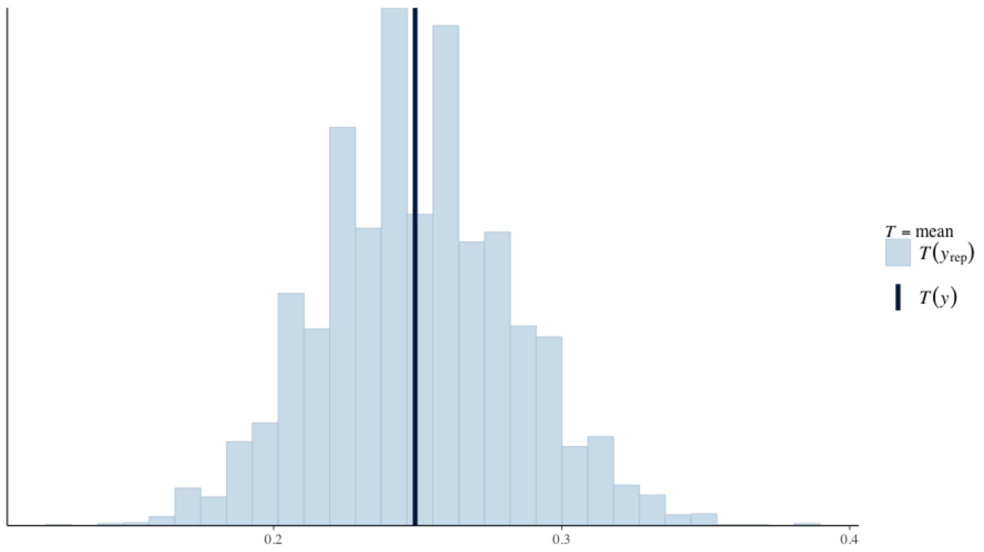


**Figure 5:** Posterior predictive distribution for $U(-\infty, \infty)$ & $N(0, 2.5)$ prior distribution.

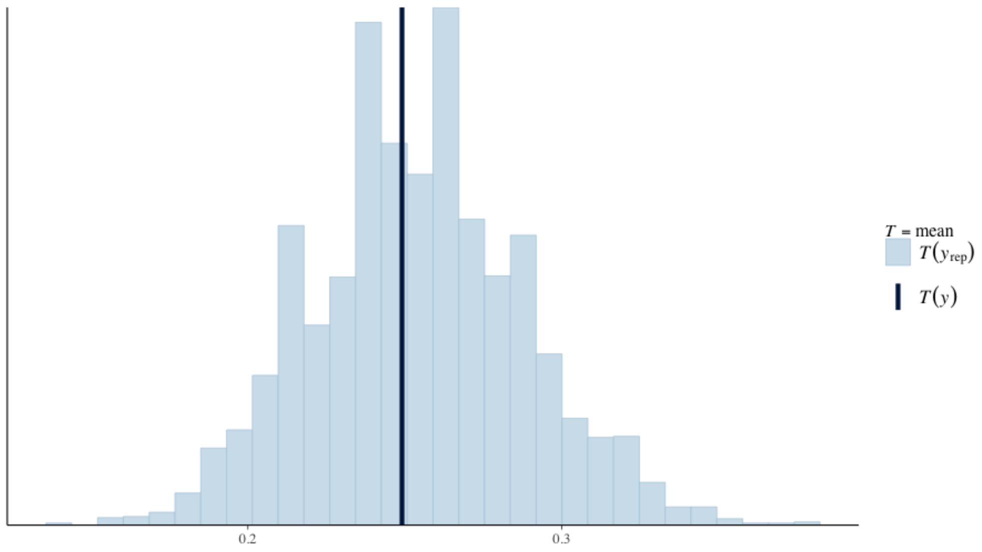


**Figure 6:** Posterior predictive distribution for $N(0, 1)$ prior distribution.

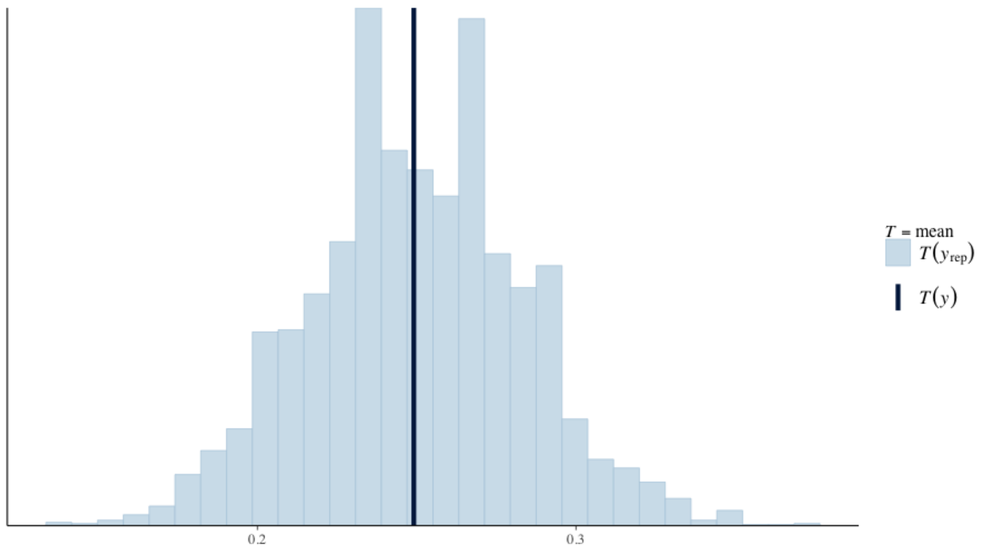


**Figure 7:** Posterior predictive distribution for $N(0, 2.5)$ prior distribution.

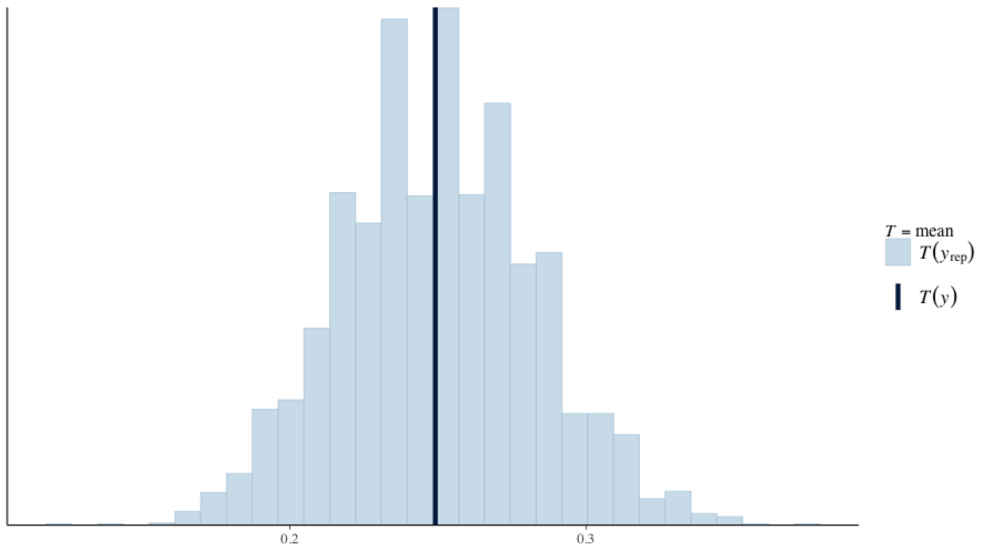


**Figure 8:** Posterior predictive distribution for $Cau(0, 10)$ & $Cau(0, 2.5)$ prior distribution.

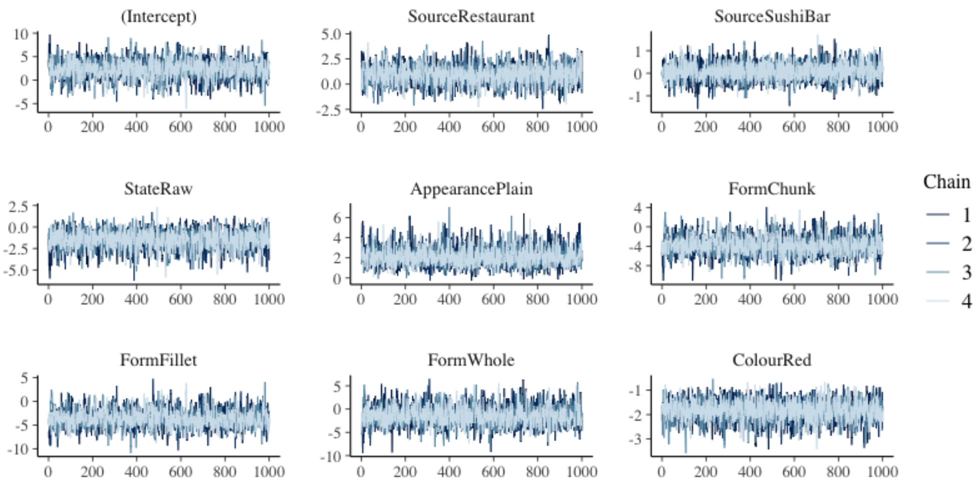


**Figure 9:** Traceplots for $U(-\infty, \infty)$ & $N(0, 2.5)$ prior distribution on both the intercept and regression coefficients, respectively.

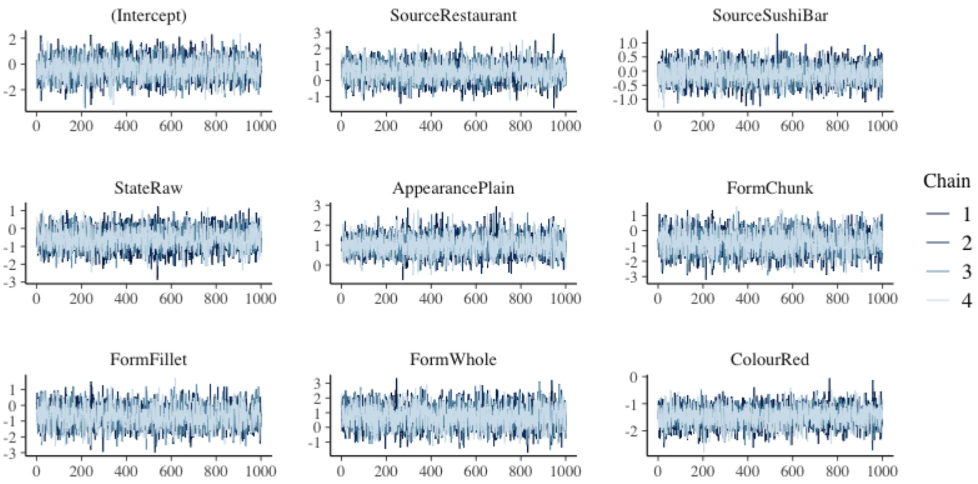


**Figure 10:** Traceplots for $N(0, 1)$ prior distribution on both the intercept and regression coefficients.

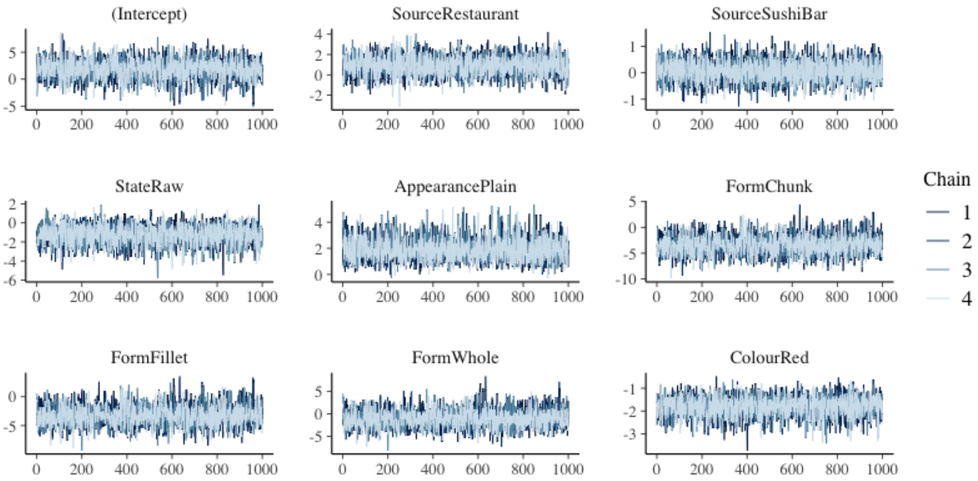


**Figure 11:** Traceplots for $N(0, 2.5)$ prior distribution on both the intercept and regression coefficients..

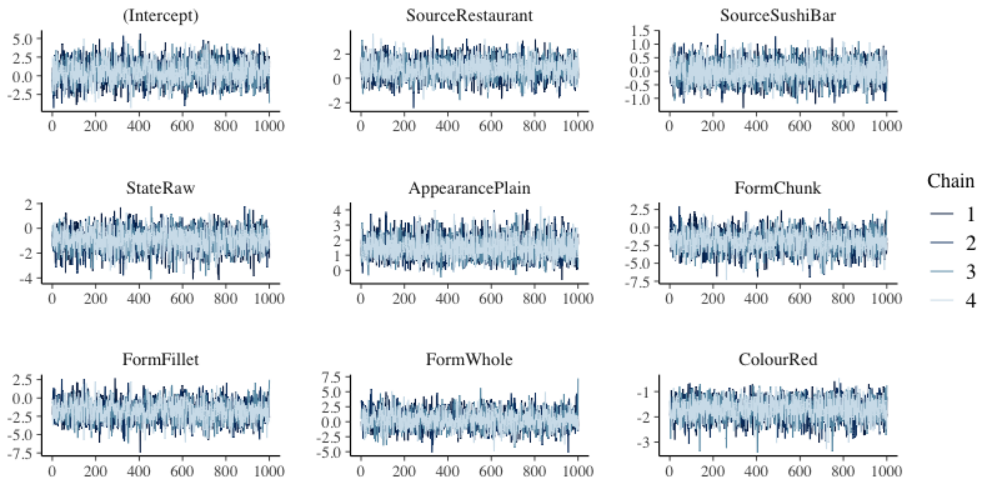


**Figure 12:** Traceplots for $Cau(0, 10)$ & $Cau(0, 2.5)$ prior distribution on both the intercept and regression coefficients, respectively.
